# The walkoff effect: cargo distribution implies motor type in bidirectional microtubule bundles

**DOI:** 10.1101/831024

**Authors:** Gleb Zhelezov, Victor Alfred, Natalia A. Bulgakova, Lyubov Chumakova

**Affiliations:** Maxwell Institute for Mathematical Sciences, School of Mathematics, University of Edinburgh, Edinburgh, UK; Department of Biomedical Science, The University of Sheffield, Sheffield, UK

## Abstract

Cells rely on molecular motors moving along an ever-shifting network of polymers (microtubules) for the targeted delivery of cell organelles to biologically-relevant locations. We present a stochastic model for a molecular motor stepping along a bidirectional bundle of microtubules, as well as a tractable analytical model. Using these models, we investigate how the preferred stepping direction of the motor (parallel or antiparallel to the microtubule growth, corresponding to kinesin and dynein motor families) quantitatively and qualitatively affects the cargo delivery. We predict which motor type is responsible for which cargo type, given the experimental distribution of cargo in the cell, and report experimental findings which support this guideline for motor classification.

## I. INTRODUCTION

Despite being highly organized, cells are in a constant state of flux, with new components being constantly produced and the old ones being destroyed [1]. They are then transported to their appropriate destinations within cells along a network of microtubules (MTs) by molecular motors. The essential cell processes relying on transport include: distribution and morphology of organelles [2–5]; segregation of chromosomes [6–8]; spindle positioning [9, 10]; and cell motility [11]. At the same time, defective transport leads to a range of conditions: neurodegenerative and cilia-dependent diseases [12–14]; and undesirable cell proliferation in cancers [15, 16].

However, the MT network, along which motors transport cargo, is constantly-changing. MTs undergo dynamic instability – an energy-intensive non-equilibrium behavior of MTs when they stochastically lengthen (grow or polymerize) or shorten (shrink or depolymerize), by the addition and loss of tubulin dimers at the polymer end [17–20], with infrequent transitions between the growth and shrinking (catastrophe and rescue). This enables MT networks to rapidly (minute-scale) adjust to cell environment, for example, entering mitosis [21, 22]. One end of the MT, the plus-end, is significantly more dynamic than the minus-end [19, 23, 24]. Parameters of MT dynamics vary depending on the cell type and the physiological state of the cell, with the growth and shrinking rates ranging between 0.03 −0.33*µm/sec* and 0.08 −0.58*µm/sec* respectively. Depolymerization rate however is consistently greater than polymerization [25–32]. Finally, this dynamics is regulated by the MT-associated proteins that bind to the plus-ends, such as the EB (end binding) family of proteins [33, 34].

There are two families of molecular motors: dyneins and kinesins. Although both walk along MTs through ATP-driven mechanochemical conformational changes, their directionality differs: dynein transports cargo towards the minus-ends, and kinesin towards the plus-ends [4, 5]. Although there are 14 families of kinesins with some moving towards minus-ends or depolymerizing MTs [4, 35], this study focuses on the paradigmal case of N-kinesins such as kinesin 1. This means that the directionality of MTs networks directly influences the motor transport. Depending on the cell type, MT networks are either unipolar, with all the MTs oriented in the same direction as in the vertebrate axons, or have mixed directionality as in vertebrate dendrites or epithelial cells [36–41]. Additionally, MTs form bundles, where several MTs are closely apposed, often connected by specialized cross-linking proteins [42–44], and unidirectional bundles facilitate intracellular transport [45, 46]. On unidirectional networks, the overall outcome of transport is intuitive: towards the minus- or plus-ends depending on the motor type, however, this outcome is non-trivial in the case of MT networks with mixed directionality.

Stable non-trivial distributions of cellular components are reliably and robustly produced on MT networks with mixed directionality, therefore there must be mechanisms in place to achieve this. Here we ask: if a distribution is established by a single motor family, what is a rule-of-thumb to predict the motor type. Answering this question requires mathematical modeling, since we aim to determine a general rule and not a particular scenario for a specific motor.

Mathematical studies of intracellular transport are numerous. The recent reviews highlight the models of stochastic motors on stochastic aligned networks are reviewed in [47], of intracellular transport on one-directional and random MT networks [48–50], and fluid dynamics effects [51]. Mathematical models of delivery via kinesin are in the system of MTs going either in the same direction (e.g. transport of neurotransmitters by kinesin in neurons), or where the MTs originate at the centrosome growing in the radial direction [52–56]. Transport along aligned networks [57], however it excluded bundled configurations. While the literature of the fundamental modeling of intracellular transport on networks is extensive, the authors are not aware of any work with goal of identifying distinct functions of each motor group on bi-directional bundled mobile MT networks.

We develop a hierarchy of models. First, our stochastic simulations incorporate both MT and motor dynamics, where we test the wide range of parameters including the growth and shrinkage of MTs, and detachment and reattachment of motors. To our surprise, different types of motors produce fundamentally different outcomes: the restricted localization, e.g. the cell boundary, requires minus-end-directed dynein motors, while the role of the plus-end-directed kinesin motors is the mixing of the components in the cell interior. The latter is due to the “walk-off effect”, since kinesin frequently changes the direction of transport due to falling off at the MT plus-ends. We validate our prediction using the concentrated distribution of E-cadherin, a key component of the cell-cell adhesion, and uniform distribution of the Golgi apparatus *in vivo*. As an example of a MT network with mixed directionality, we use the apical MT network in the elongated cells of the *Drosophila* embryonic epidermis [58]. Finally, we develop a probabilistic toy model of transport on a bundle, which reveals the underlying difference in the motor behavior. The differential equations for the delivery time to the cell boundary are drastically different for the two motors (regular for dynein, and singular for kinesin), making a reliable targeted delivery by kinesin impossible.

Altogether we demonstrate that on MT networks with mixed directionality in the absence of additional mechanisms the rule-of-thumb to determine the motor type, given the distribution of cellular components, is: if a single motor family is involved, the uniform distribution of cellular components is achieved by kinesin, while the localized distribution - by dynein.

## II. STOCHASTIC SIMULATIONS SHOW THAT DYNEIN RELIABLY DELIVERS CARGO TO THE CELL BOUNDARY

To test *in silico* the qualitative difference between the dynamics of kinesin and dynein on a MT bundle, we combine two stochastic models - of a MT and of a motor.

The 1d MT bundle consists of stochastic 1d MTs anchored at cell walls. Each MT is modelled by a finite length Markov chain (Fig.1a) to account for the dynamic instability (following [58–60]). Here the MT of length *n* dimers can be in either polymerizing, depolymerizing, or the minus-end state (*P*_*n*_, *D*_*n*_, or *D*_0_), where the maximum MT length is the length of the bundle. MT dynamic instability rates are: polymerization -*α*, depolymerization -*β*, rescue -*α*′, and catastrophe -*β*′.

**FIG. 1.**
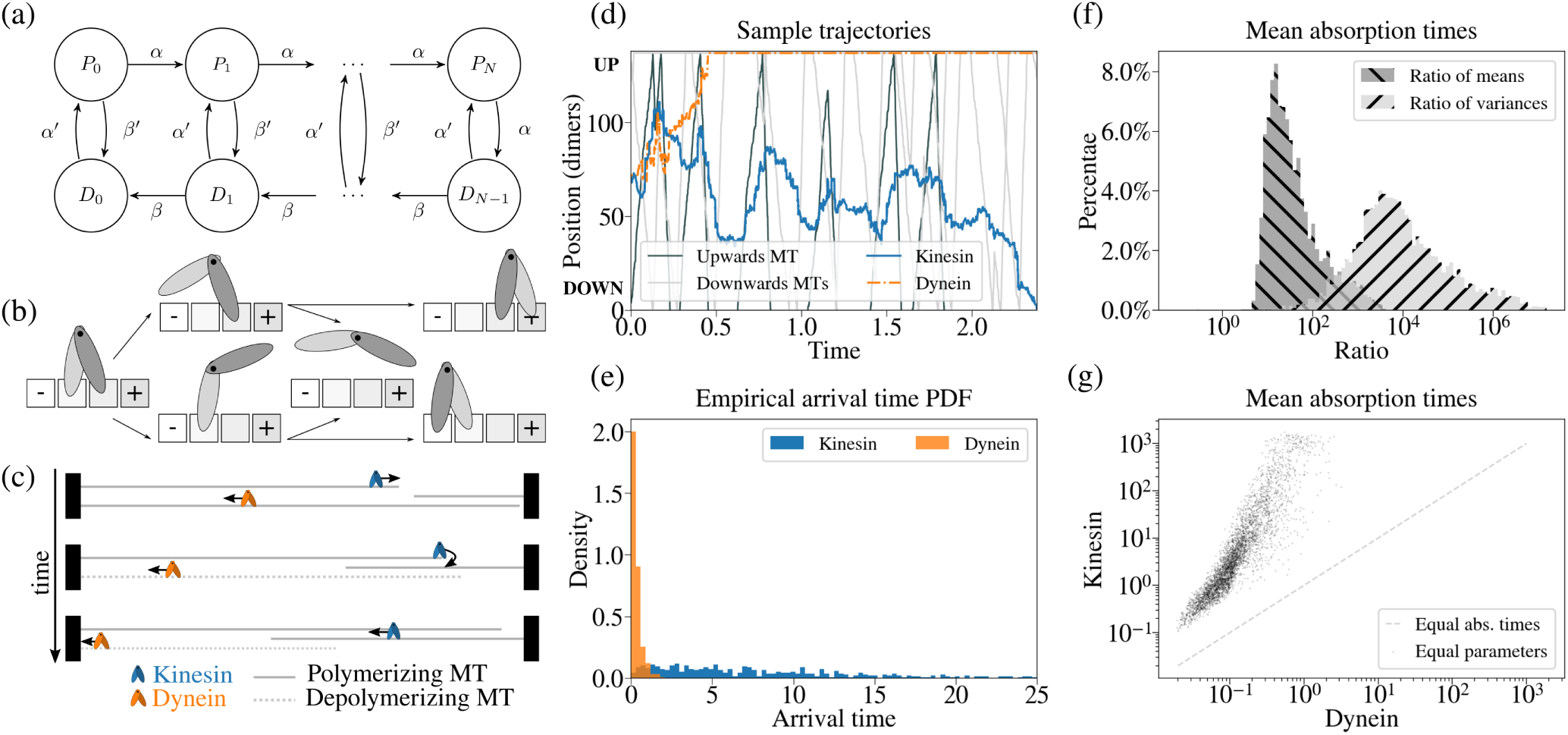
Stochastic simulations of kinesin (*blue*) and dynein (*orange*) on a MT bundle. (a) Markov chain MT model. *P* and *D* are polymerizing and dipolymerizing states of a MT, (·)_*n*_ indicates that the MT length is *n* dimers, with the maximum length *N*. The rates are *α* - of polymerization, *β* - depolymerization, *α*′ - rescue, and *β*′ - catastrophe. (b) Model of molecular motor on a MT. *±*-indicates the directionality of a MT (towards plus- or minus-ends). *Left-to-right*: when both motor heads are attached to a MT, one can detach, then the second head either attaches or detaches, leading to the motor either finishing the step or falling off a MT. (c) Three consecutive instances of motors on a bundle. MTs (polymerizing -*solid*, depolymerizing -*dashed*) are anchored at the cell boundaries (*black rectangles*). Kinesin moves towards the MT plus-end, outruns it (*top*) walking off the MT (*middle*), and reverses its direction by attaching to an oppositely directed MT (*bottom*). Dynein, moving wards MT minus-end (*top*), outruns the MT depolymerization (*middle, bottom*). (d) Typical kinesin and dynein trajectories on a MT bundle (*vertical*) as a function of time. *Grey* trajectories are MT plus-ends (1 MT growing upwards -*solid*, 2 MTs downwards -*dashed*). (e) Empirical distributions of the motor arrival to the cell boundary for kinesin and dynein. (f) Histograms for all the stochastic simulations of the ratios (dynein to kinesin) of the mean arrival time (*dashed negative slope*) and its variance (*dashed positive slope*). (g) Scatter plot of dynein and kinesin mean arrival times. For every parameter set we ran 200 simulations to compute the mean and the variance. The details of the sampling are in the Appendix E.

We model both motor types on a bundle via a hinged walker model (Fig.1b, [59]), Since both types can occasionally step backwards [61–63], we term the next dimer to them “front” and “back”, depending on the typical directionality of the motor (e.g. for dynein “front” is towards the minus-end, while for kinesin it is “back”). The motor can transition between the following states: a motor with two attached heads can detach one of them (front or back); at the next time-step, it can either enter a diffusive state by detaching the bound head, or remain on the MT by attaching the free head to the front or to the back. The latter either completes a step, or returns the motor to the original position. When the motor reaches the MT plus-end, it falls off. Although some kinesins can track MTs plus-ends, the residence time of the classical cargo carrier kinesin-1 at the plus-ends is low [64]. When the motor reaches the cell boundary, we term that the motor has delivered the cargo.

Combining the two stochastic models introduces a 14-dimensional parameter space, where the parameters are the transition rates in both the MT and walker models, and the bundle length. We ran stochastic simulations for a sampled range of parameters (see Appendix E), ensuring that the biological aspects of this dynamics remain valid. In particular, we preserved the generic scale-separation for velocities: kinesin outruns MT polymerisation, while dynein outruns depolymerisation. Both motors reach velocities of over 1*µm/sec* [65–69], which are around 5-fold greater than the polymerisation and depolymerisation rates of MTs. Furthermore, both motors achieve long-range transport over 1*µm* (a significant fraction of the cell length) due to their high processivity, suggesting low fall-off frequency along the MT lattice [62, 70–72]. The consequences of this scale separation are demonstrated on the Fig.1c. On a MT bundle, in the three consecutive time-steps, dynein outruns MT depolymerization and delivers cargo to the left cell boundary, while kinesin *walks-off* a MT: it falls off upon reaching a plus-end, and re-attaches to another MT. In the simulations, the motor in the diffusive state has equal probability to attach to any MT in the bundle, including the one it walked off.

The simulations revealed drastically different behavior of the two motor types (Fig.1d-g). Sample trajectories in Fig.1d show that while dynein reaches the cell boundary in a fairly direct manner, kinesin spends a large fraction of time in the cell interior, switching directions multiple times before reaching the boundary. This makes the arrival times for dynein small and have a low variance, while the the variance for the kinesin arrival time is very large, perhaps limited by the extent of our sampling (Fig.1e). To compare the two motor behaviors, we set kinesin and dynein to differ only in the preferred stepping direction. Fig.1f,g show that in this case kinesin not only arrives to the boundary much later than dynein, but the arrival time variance is order of magnitudes larger.

The stochastic simulations therefore suggest that the two motors have different functions in cells with bidirectional bundled MTs. In particular, dynein is efficient for targeted delivery (e.g. the cell boundary), while kinesin could be used for an uniform re-distribution of the cellular components.

## III. EXPERIMENTAL EVIDENCE FOR DISTINCT MOTOR FUNCTIONS

### A. MT bundles are bidirectionally oriented in *Drosophila* embryonic epidermis

To validate the predicted roles of kinesins and dyneins in transport on MT networks with mixed directionality we turned to epithelial cells in the *Drosophila* embryonic epidermis. There, the apical MTs grow from the sites of cell-cell adhesion on the cell boundaries and are restricted to a thin 1 *µm* layer below the cell apical surface [26, 38, 60]. To investigate the directions of MT growth, we examined live embryos expressing the GFP-tagged End Binding protein 1 (EB1-GFP) at the stage 15 of the embryonic development (Fig. 2a). We tracked each EB1-GFP comet and determined the direction of growth relative to the dorsal-ventral embryonic axis, which the cell major axes are aligned with [38]. Consistent with the findings that the cell geometry aligns the MT network with the cell major axis in elongated cells [26, 73], the MT growth was aligned with the direction of the tissue (Fig. 2a and Movie S1). Due to the lack of cell border information, we estimated the cell dimensions by taking into account the developmental stage and image acquisition parameters to tile the tissue into cell-scale areas (Fig. 2b). To ensure that there was no bias due to random choice of tile position, we also tiled the same tissue with a half cell-size offset (Fig. 2b). The ratio between MT growth events in positive and negative directions relatively to the embryonic axis did not differ from 1 for individual tiles in each embryo (examples are in (Fig. 2c), and on average per tiling in individual embryos (Fig. 2d). Therefore, at both the tissue and cell levels, MT growth events were distributed equally across opposite directions relative to the main axis of the cell, which makes the embryonic epidermis a perfect system for testing the model predictions about the roles of kinesin and dynein in transport along MT networks with mixed directionality.

**FIG. 2.**
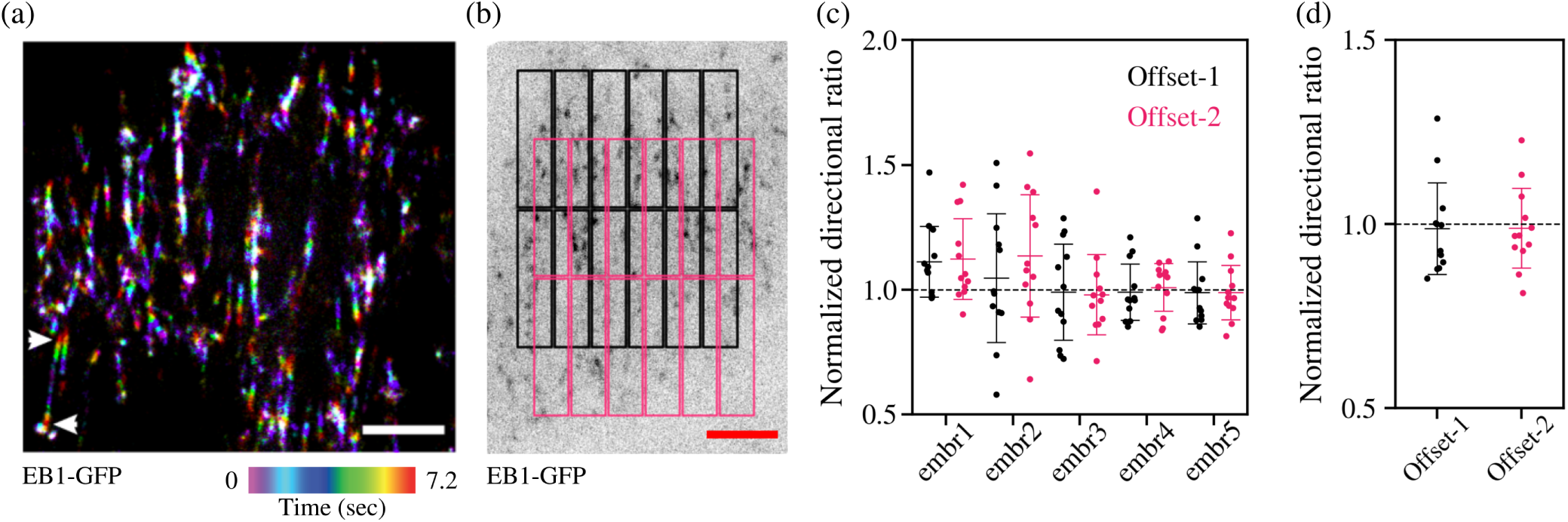
Apical MTs have mixed directionality in the *Drosophila* embryonic epidermis. (a) Example of MT growth in the *Drosophila* embryonic epidermis. The MT growth was visualized by EB1-GFP and the time series was projected in the displayed color-code. Arrowheads indicate an example of two MT growing in opposite directions next to each other. The scale bar is 5 *µm*. (b) Example of tiling used for confirming mixed-directionality of the MT network (*black*, offset-1) and same tiling with an offset (*magenta*, offset-2). (c) Ratios of MT growth events in the positive and negative directions relative to the dorsal-ventral embryonic axis in each tile without or with offset in 5 different embryos. Each dot represents an individual tile. (d) Average ratios of MT growth events in positive and negative directions relative to the dorsal-ventral embryonic axis in a tile without or with the offset. Each dot represents an individual embryo.

### B. Dynein, but not kinesin, affects directional delivery of E-cadherin

To test our mathematical predictions, we analysed the effect of depleting kinesin and dynein motors on transport in the *Drosophila* embryonic epidermis. E-cadherin, a key cell-cell adhesion molecule, is transported along MTs towards the cell boundaries [26, 74]. Alignment of MTs with the cell major axis leads to asymmetry in E-cadherin distribution: the levels of E-cadherin at the short cell borders perpendicular to MT direction (40-90° to the cell major axis) are almost 2-fold larger than those on the cell long borders (0-10° to the cell major axis, Fig. 3a-c, [26]). This asymmetry provides a reliable read-out of the MT-dependent E-cadherin delivery [26].

**FIG. 3.**
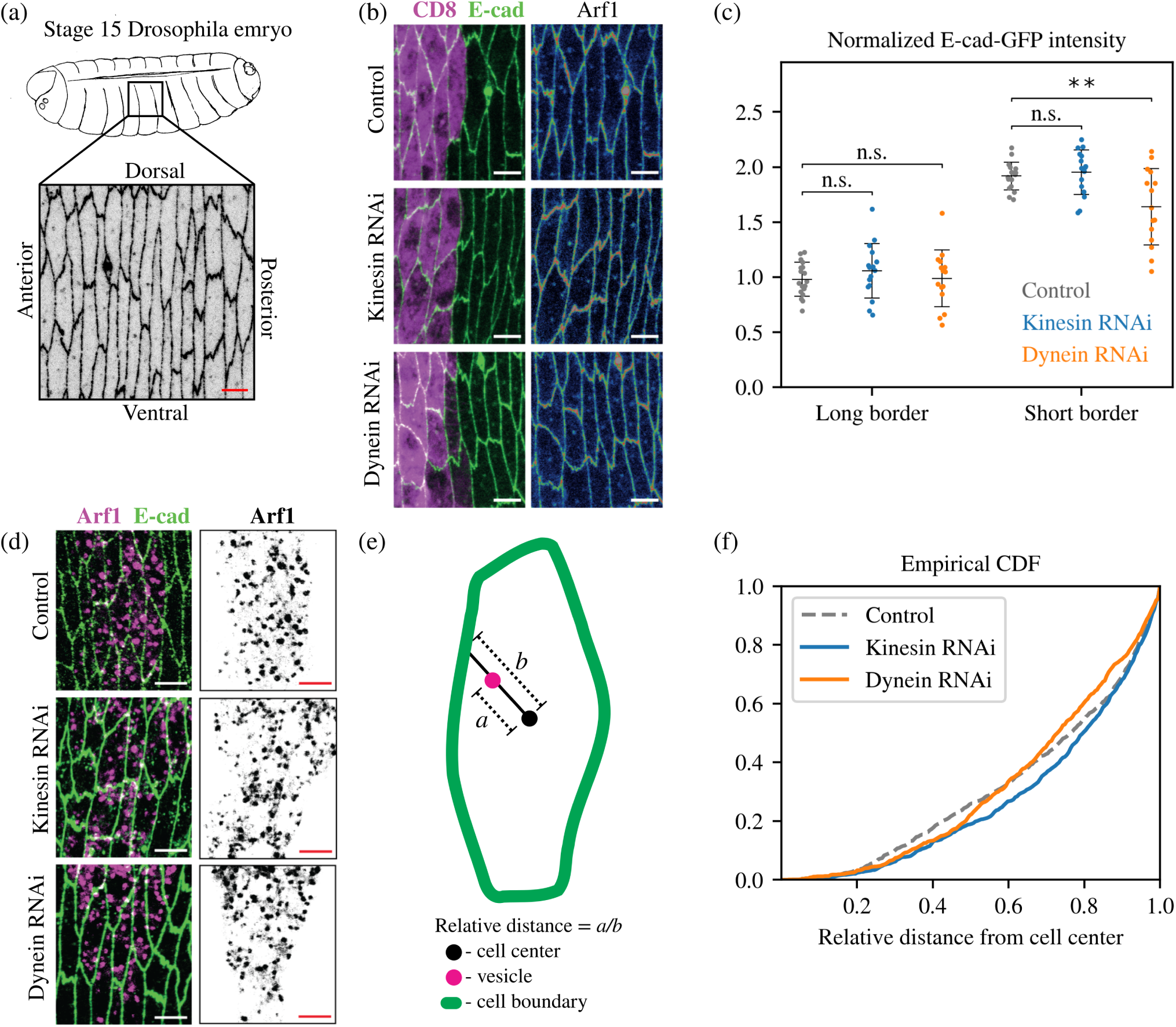
Roles of kinesin and dynein in intracellular transport in the *Drosophila* embryonic epidermis. (a) Schematic diagram of the *Drosophila* embryo at the stage 15 of the embryonic development with an example of the surface view of the epidermis, where the cell boundaries are visualized with GFP-tagged E-cadherin. The scale bar is 5 *µm*. (b) Examples of E-cadherin distribution in the control cells (*top*), and the cells where kinesin (*middle*) or dynein (*bottom*) were depleted with RNAi. E-cadherin visualizes cell boundaries and its intensity was used for quantification (*green, left*; *rainbow intensity profile, right*). Cells expressing RNAi are visualized by CD8-Cherry (*magenta, left*). The scale bar is 5 *µm*. (c) E-cadherin-GFP (E-cad-GFP) average levels at the long borders and short borders (0-10° and 40-09° to the dorsal-ventral embryonic axis) in control and cells expressing RNAi against kinesin or dynein. ** - p = 0.004. (d) The Golgi apparatus visualized with Arf1 protein tagged with GFP (*magenta, left*; *grey, right*) in control (*top*) and cells where kinesin (*middle*) or dynein (*bottom*) were depleted with RNAi. Cell outlines are visualized by antibody staining of E-cadherin (*green, left*). The scale bar is 5 *µm*. (e) The diagram of quantifying distribution of the Golgi apparatus: the distance between the Arf1-positive spot and the cell center was divided by the distance between the cell center and the boundary on the line drawn through the center and the Arf1-positive spot. The resulting ratios were used to produce the cumulative distributions in (f). (f) Cumulative distance distributions of Arf1-positive spots (the Golgi apparatus) in the control and in the cells where kinesin or dynein were downregulated with RNAi.

Using RNA interference (RNAi) against the dynein heavy chain (*Dhc*), kinesin heavy chain (*Khc*), as well as several other genes encoding motor heavy chains (Fig. 3a-c and Fig. 5a), we depleted the cellular levels of these motors in stripes of cells using the *engrailed*::GAL4 driver expressed in the posterior half of each embryonic segment. This provided us with a side-by-side comparison of perturbed and wild type (control) cells within each embryo. Cells depleted of kinesin or dynein showed similar levels of E-cadherin at the long borders relative to control cells (0.979 ±0.154, 1.057 ±0.246, 0.988 ±0.257 respectively, Fig. 3b,c). While E-cadherin levels at the short borders were similar between control and kinesin-depleted cells (1.918 ±0.127 vs. 1.953 ±0.203, *p*-value = 0.5544. Fig. 3b,c), dynein-depleted cells exhibited significantly reduced levels of E-cadherin at the short borders relative to the control cells (1.918 ±0.127 vs. 1.638 ±0.346, *p*-value = 0.004, Fig. 3b,c). This was not due to changes in cell shape, as neither kinesin nor dynein affected cell area or eccentricity (Fig. 5b,c). changes in E-cadherin levels at either type of borders were observed for other RNAi (Fig. 5a). This demonstrates that dynein, rather than kinesin, facilitates the MT-mediated delivery of E-cadherin to the cell borders, which is consistent with the prediction of our stochastic model that dynein rather than kinesin is efficient in concentrating cellular components in specific locations.

**FIG. 4.**
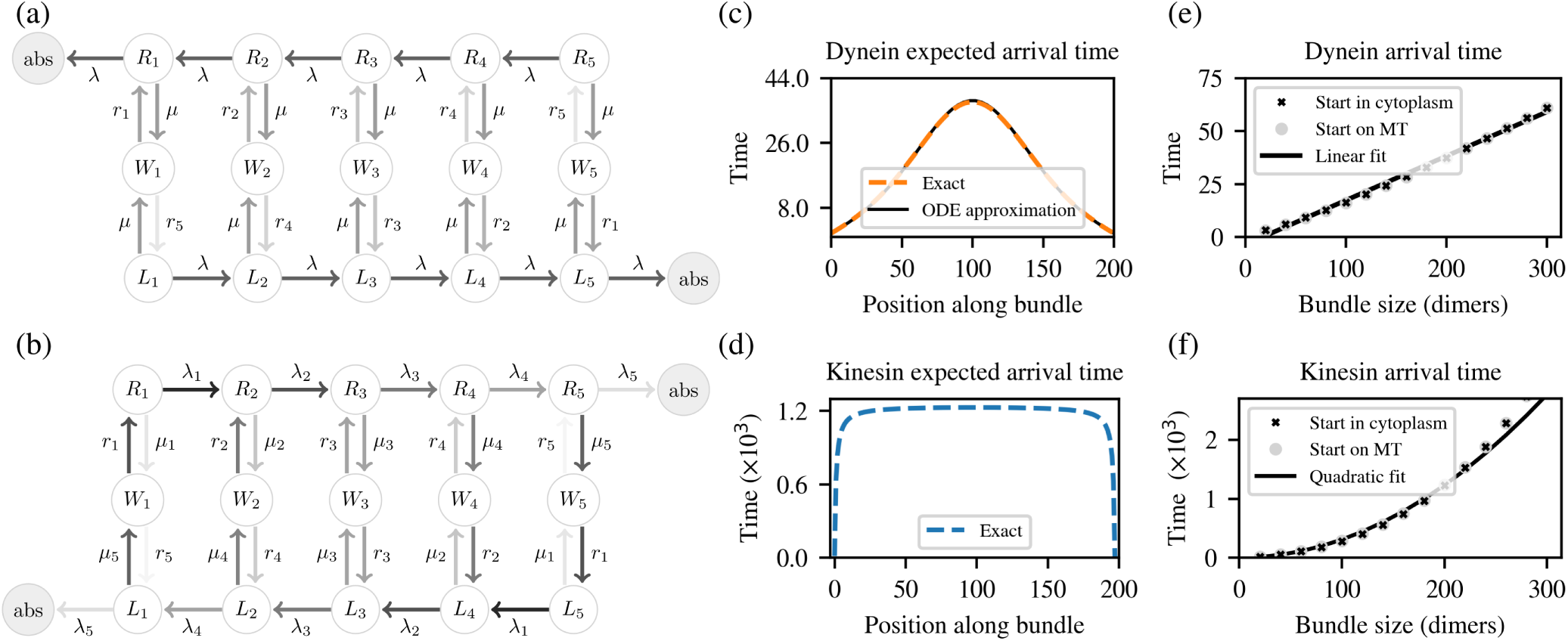
Analytical model highlights that the origin of the walk-off effect. (a,b) Molecular motor state diagram for dynein (a) and kinesin. Motor states at the *i*th position along the dimer are *R*_*i*_ (*top*) along the right-directed MT, *L*_*i*_ (*bottom*) along the left-directed MT, *D*_*i*_ (*middle*) is the waiting lattice, *abs* is the absorbing state, when the motor has arrived to the cell boundary. The rates are *λ* - walking, *µ* - detachment, *r* - reattachment. The index (·)_*i*_ indicates the dependence of the rate on the position of motor along the bundle. (c,d) Arrival time as a function of the initial position on a bundle for dynein (c) and kinesin (d). (e,f) The maximum arrival time and a polynomial fit as a function of the bundle length *N*: (e) arrival times of dynein from the cytoplasm, and the fit *y* = 0.209*N*; (f) arrival times of kinesin from the cytoplasm on the right-going MT, and the fit *time* = 0.031*N* ^2^.

**FIG. 5.**
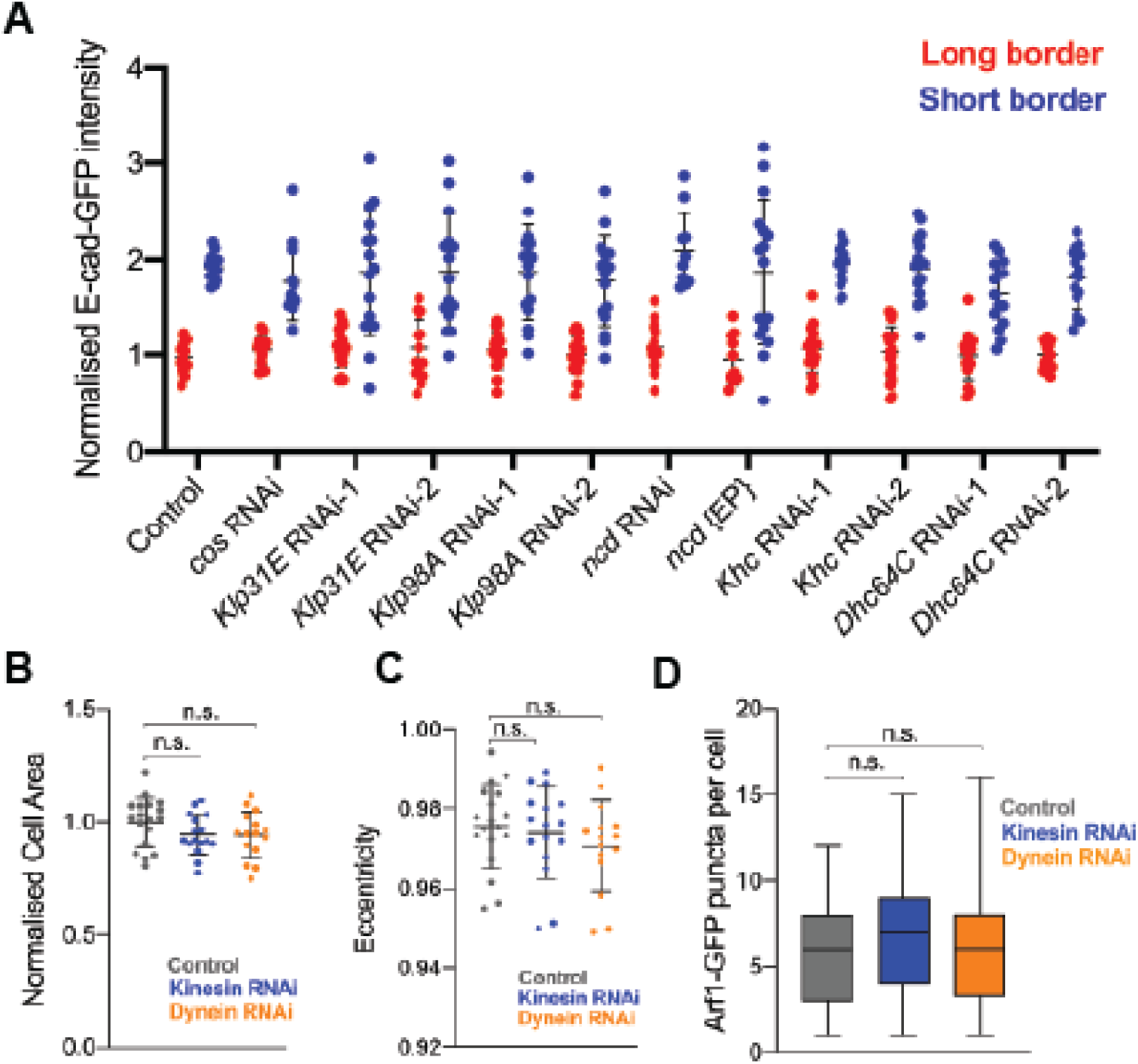
Effects of motor depletion on E-cadherin levels and cell shape. (a) E-cadherin-GFP (E-cad-GFP) average levels at the long borders (0-10° to tissue axis) and short borders (40-09° to tissue axis) in control and cells expressing RNAi against genes encoding heavy chains of different motors. The only changes were observed for RNAi against dynein (*Dhc64C*), also shown in Fig. 3b,c. (b-c) Average cell shape in control and embryos expressing RNAi against kinesin or dynein. Average cell area (b) and eccentricity (c) are shown with each dot depicting a single embryo. (d) Average number of Arf1-GFP (Golgi) puncta per cell. A minimum of 120 cells were analysed in each group. ns - not significant.

### C. Kinesin regulates distribution of the Golgi apparatus in the *Drosophila* embryonic epidermis

To test the second prediction that the uniform cargo distribution might be more favoured by kinesin activity rather than dynein, we selected the Golgi apparatus, whose distribution is dependant on MTs [75–77]. We used the GFP-tagged Arf1 protein to visualized the Golgi apparatus [78]. In epidermal cells, Golgi appears as multiple spots of Arf1 in the plane of cell-cell adhesion and MTs (Fig. 3d), consistent with fragmentation of Golgi, also reported in other differentiated cells [79]. We examined distributions of the Arf1-positive Golgi spots in control cells and those where kinesin or dynein were depleted with RNAi (Fig. 3d). To do so, we measured the relative distance of these spots from the cell center (Fig. 3e): namely, we calculated the ratio of the center-spot to center-boundary distances.

We observed, that kinesin but not dynein depletion resulted in an increased accumulation of the Golgi apparatus near cell boundaries (p=0.01 and p=0.21, Fig. 3f), as apparent from the shift of the cumulative distribution in kinesin-depleted cells to the right. This indicates that when kinesin is downregulated, the cells are less efficient in uniformly distributing the Golgi apparatus, and it accumulates in the proximity of the cell boundaries instead.

Altogether, these two experimental examples, E-cadherin delivery and Golgi positioning, support the prediction of our stochastic model that the roles of kinesin and dynein in transport along MT networks with mixed directionality are drastically different.

## IV. WALK-OFF EFFECT IS PRIMARILY RESPONSIBLE FOR THE DELIVERY DIFFERENCE BETWEEN THE MOTOR TYPES

If the fundamental difference between the outcomes of kinesin and dynein transport were only due to the walk-off effect, its causes would be independent of the motor diffusion in the cytoplasm and along MTs, since those vary between biological systems. We therefore explore simplified models, which allow for tractable analytical solutions. We introduce a 1d MT bundle with the maximum MT length of *N* dimers, and two same-size families of MTs anchored at the opposite cell boundaries (Fig.4a,b). Then the motor at the *i*th dimer location along the bundle can either attach to a MT and be in a *L*_*i*_ or *R*_*i*_ state depending if the MT is of the left- or right-directed families, or be in the cytoplasm in the waiting state *W*_*i*_. We will use the arrival time to the cell boundary (the *abs* state in Fig.4a,b) as a function of the initial motor position *i* in the cytoplasm to compare the transport efficiency by the motors.

On a single long MT, the motor walking rate is *λ*, and the fall-off –*µ*. For a motor in a waiting state next to this MT, the rate of reattachment is *r*. However, when a motor is in the vicinity of or on a MT bundle, these rates of switching between the *L*_*i*_, *W*_*i*_ and *R*_*i*_ states depend on the motor position, as we describe below. From the cytoplasm, the probability of reattachment to a MT is proportional to the local MT probability density. For the full Markov Chain MT model in Fig.1a, the probability of MT being of a particular length is geometric (see Appendix F for details). Here we simplify it to be a linear function, since we are looking for tractable analytical solutions that capture the walk-off effect. The reattachment rates for both motor types to the right-directed MT are therefore

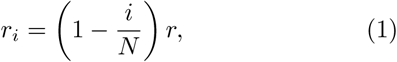

and to the left-directed MTs, with the reversed indices *i* ↔*N* −*i*. Dynein stepping rate *λ* and fall-off rate *µ* are taken to be constant, since dynein outruns the MT depolymerization [27, 28, 67, 69]. However, for kinesin we have to account for the walk-off effect. When attached to the *i*th dimer on a right-going MT, it can only step forward if the MT is longer than *i* dimers. Therefore, its walking rate decreases with *i*

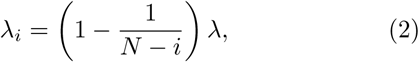

and the fall-off rate is increasing with *i*

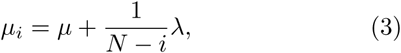

where the indices are reversed, *i* ↔*N* −*i*, for the walking and detachment rates along the left-directed MT.

Then, the dynein expected absorption times (Fig.4a), denoted by 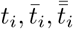 for a motor starting in the *R*_*i*_, *W*_*i*_, or *L*_*i*_ respective states, satisfy

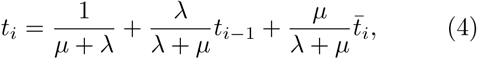

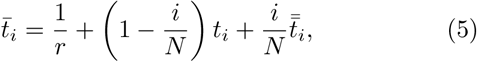

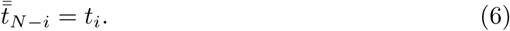

This reduces to the following equation for the expected times *t*_*i*_ starting on the right-directed MTs

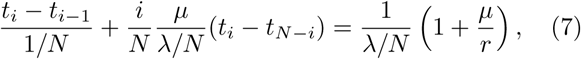

which we approximate by the first order ODE

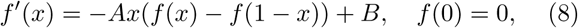

where

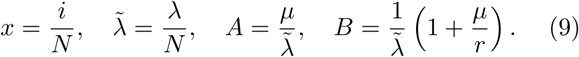

Here *x* is the rescaled motor position along the bundle, 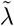 is the walking speed. The ratio of the fall-off rate to the walking speed (*A*) determines the solution shape, and a combination of the walking speed and the ratio of the fall-off to the reattachment rates (*B*) determines the solution amplitude. This ODE is unusual, as its right-hand-side depends on the odd part *p*(*x*) = *f* (*x*) −*f* (1 −*x*) of *f* (*x*) around the middle of the bundle *x* = 1*/*2.

Rewriting Eq.8 as an ODE for *p*(*x*) and solving for *f* (*x*) and *p*(*x*), we obtain

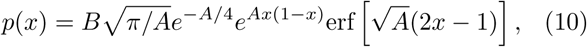

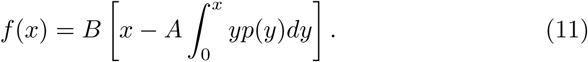

The arrival time from the cytoplasm (state *W*_*i*_) is then

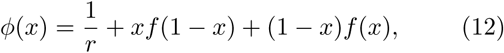

where the solution amplitude is largely determined by *B*, namely the motor speed and the ratio of the fall-off to the reattachment rates, while the shape of the graph is determined by *A*, namely the ratio of the fall-off rate to the walking speed.

In contrast, the similar ODE for kinesin is singular. Consider the arrival times from the *i*th dimer, as before,

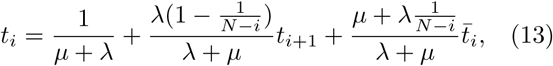

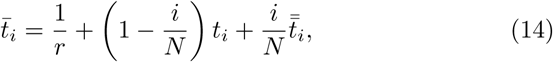

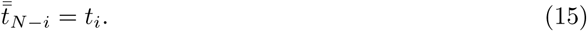

Then for the arrival times *t*_*i*_ for a motor starting on a right-directed MT satisfy

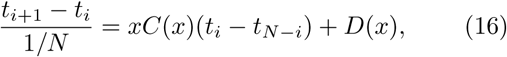

where *x* = *i/N* is again the relative position of the motor along the dimer, *ϵ* = 1*/N*, and

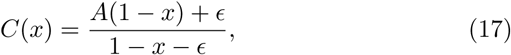

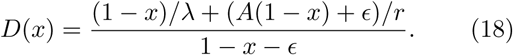

Eq.16 has to be approximated by a second order ODE

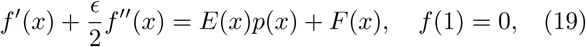

where

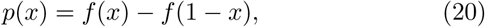

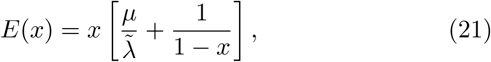

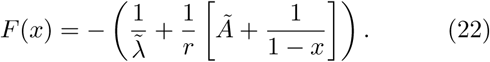

Eq.19 has a boundary layer at *x* = 1, where the outer solution is given outside the *ϵ* - thin boundary layer at *x* = 1 by solving Eq.19 for *ϵ* = 0, and the inner solution inside the boundary layer grows like *f* (*x*) ∼(*x/ϵ*)^2^. This scaling was obtained by changing variables for the inner solution *Y* (*X*) = *f* (*x*) as a function of *X* = (1 −*x*)*/ ϵ*. Then *Y* satisfies *XY* ^″^ + *XY* ′ −*Y* ′ + *Y C* = 0, where *C* is a constant, *Y* (0) = 0, and has the leading order behaviour near *X* = 0 can be shown to be *Y* (*x*) ∼ *X*^2^ using Frobenius method.

The arrival time from the cytoplasm as a function of the starting dimer position (*ϕ*(*x*), Eq.12) indeed has two boundary layers at the first and the last dimer (Fig.4d). We confirm that its maximum is closely determined by the maximum arrival time starting on the right-directed MT (*f* (*x*)), and indeed scales as a square of the bundle length in dimers (Fig.4f). This quadratic growth is the lower bound, since it only reflects the increase of *f* (*x*) in the boundary layer; the further increase can be seen for *N >* 200 (Fig.4f). In contrast, the maximum arrival time for dynein grows linearly with the bundle size (Fig.4e).

This confirms that the distinct behaviour of the two motor families is due to the walk-off effect, which introduces singularities in the differential equations for kinesin. Additional effects, e.g. diffusion in the cytoplasm and along the MTs, would regularize it, but the gap in the arrival times of kinesin and dynein would remain large. Furthermore, the more realistic MT distribution in the bundle decays faster than linear away from the MT minus ends (see Appendix F), and hence using it in Eq.1-3 would strengthen the walk-off effect, as kinesin would be more likely to walk-off MTs further from the target cell boundary.

## V. DISCUSSION

Here we provide a rule-of-thumb in cell biology of how to determine the motor type in the systems with MT networks of mixed directionality from the steady state distributions of cellular components delivered by these motors. The rule is: if the distribution is uniform – the components are transported by kinesin, if concentrated in specific locations – by dynein. We further suggest that the two motor families have distinct biological roles in this system: the role of kinesin is mixing, while that of dynein is the targeted delivery.

While targeted delivery is vital for the correct cell function, there is an emerging evidence that the cellular cytoplasm undergo mixing by fluid flows [80]. These flows contribute to organization of microtubule network, and consistently with our model require kinesin function [81]. We further suggest that such constant mixing of some of cellular components, might be crucial for efficient intracellular trafficking, when different vesicle compartments need to fuse during the maturation process. For example, late endosomes fuse with lysosomes for protein degradation [82]. Similarly, endosomes fuse with the Golgi apparatus, studied here, for protein recycling [83]. The mixing in the cytoplasm by kinesin, is likely to enable the Golgi to cover larger area and facilitate meeting and thus fusing with other compartments.

Our work highlights the importance of MT bungling, as it prevents the passive diffusion of a detached motor into the cell interior, thus aiding the ballistic motion of the motor (though not necessary while preserving the direction). The sequence of models in our study was chosen to highlight the walk-off effect, hence we omitted such processes as the tug-of-war (motor co-operation or retardation) or role of flows created by the motors. However, we suggest that the walk-off effect is the main driver of the difference between the two motor functions. These additional processes open avenues for future studies of transport by specific motors, considering individual differences in behaviours, e.g. kinesin-1 vs kinesin-13 [64], as well as differential post-translational modifications of MTs [84]. Finally, while our modeling results were analyzed on the example of a 1d bundle, we interpret it as a 1d projection of a pseudo-2d system, such as in our *in vivo* model, suggesting that the rule-of-thumb is applicable for a broad range of MT networks of mixed directionality.

Here, we highlight the importance of the generic walk-off effect of kinesin, supporting this principle by the results of our stochastic simulations and *in vivo* experiments. We further hypothesise that in biological systems, the additional differences in the the exact details/statistics of the delivery would provide flexibility, which is required for an organism to produce a multitude of cell types with different shapes and functions.

## Appendix A Fly stocks

Flies were cultured at 25 °C on a 12h light: 12h dark cycle in vials containing fresh standard medium. The following fly stocks were used in this study: *w*^*1118*^ (Bloomington 3605), *UAS*::CD8-mCherry (BL 27392), *engrailed*::GAL4 (BL 30564), *shg*::E-cad-GFP (BL 60584), *UAS*::Dhc64C-RNAi (Dynein heavy chain 64C, BL 36583, 36698), *UAS*::Khc-RNAi (Kinesin heavy chain, BL 35409, 35770), EB1-GFP ([26]), *UAS*::Arf1-GFP (gift from T. Harris).

## Appendix B Immunostaining

Fly embryos were collected at 3-hour time intervals, allowed to develop until stage 15, and dechorionated in 75% sodium hypochlorite (bleach, Invitrogen) in water for 4 min. Embryos were washed repeatedly in deionized water to remove excess bleach, fixed with a 1:1 solution of 4% formaldehyde in PBS:heptane for 20 min at room temperature, and devitellinized by vigorous agitation for 30 s in 1:1 methanol:heptane. Following devitellinization, embryos were washed 3 times in methanol and then three times in PBST (PBS containing 0.05% Triton-X). Embryos were either imaged directly or stained with antibodies. 1% normal goat serum (NGS) in PBST was used to block embryos for 1 hour at room temperature followed by incubation with primary anti-E-cadherin antibody (DCAD2, DSHB, 1:100) overnight at 4 °C. This was followed by three washes in PBST, and incubation with anti-rat Alexa Fluor 568-conjugated secondary antibody overnight at 4 °C. Embryos were again washed three times, incubated for 1 hour in Vectashield (Vector Laboratories) and mounted on glass microscope slides.

## Appendix C Microscopy

For fixed samples, images were acquired on an upright Olympus FV1000 confocal microscope with a 60x/1.40 NA oil immersion objective. All experiments were performed on the dorsolateral epidermis of stage 15 embryos. 16-bit 1024×768 pixel XY-images were taken at magnifications of either 0.058 µm/pixel (for Arf1-GFP experiments) or 0.078 µm/pixel (for E-cad intensity experiments). Six z-axis sections per embryo were obtained at 0.38 µm spacing.

For EB1-GFP live imaging, embryos were dechorionated 50% bleach, washed in water, and embedded in halocarbon oil 27 (Sigma-Aldrich) on the surface of a glass dish. Image acquisition was made with an inverted microscope (Eclipse Ti-E; Nikon), equipped with a CFI Apochromat total internal reflection fluorescence (TIRF) 100 1.49 NA oil objective lens (Nikon) and a motorized confocal head (CSU-X1-A1; Yokogawa). 16-bit images were projected onto the CCD chip at a magnification of 0.045 /pixel at a frame rate of 1.67 per second.

## Appendix D Image and data analysis

Maximum and average projections of images were performed using Fiji. Maximum intensity projections were used to cell masks while the intensity quantification was done on average projections. Cell borders were outlined using the TissueAnalyzer plug-in in Fiji.

### Membrane E-cad intensity

Quantification was performed with a custom MATLAB script, as previously described [73].

### Arf1-GFP puncta distribution

The relative distance of Arf1-GFP puncta from the cell border was automatically determined on a cell-by-cell basis using a script we developed in MATLAB. Briefly, maximum projections of the Arf1GFP signal were binarised by applying a global threshold using Otsu′s method, and then de-noised using a median filter. Then, cell outlines obtained from TissueAnalyzer were used to identify each cell as an individual object. Next, binary Arf1-GFP signal within each cell was filtered using the cell outline as a mask and the centroid position of each object was determined. We fitted a straight line through each Arf1-GFP object and the cell mask centroid and obtained the coordinates at which this line intersects with the cell border. We calculated the relative distance as the ratio of the Euclidean distance between the Arf1-GFP object and cell centroid *a*, and the Euclidean distance between the cell centroid and border closest to the Arf1GFP object *b* (Figure 3E). A relative distance of 1 represents an Arf1-GFP object located on the cell border while a value of 0 means an object positioned in the centre of the cell.

### EB1-GFP MT dynamics

The time-series with EB1-GFP signal in the dorsolateral epidermis did not contain outlines of individual cells for technical reasons. To estimate MT dynamics on a cell-by-cell basis, we generated cell masks in Fiji (http://fiji.sc/) based by tiling rectangles corresponding to individual cell dimensions onto the image. Individual EB1-GFP comets were identified and tracked using the TrackMate Fiji plug-in. Information on the direction and length of tracks were determined using custom scripts in MATLAB and R. To avoid bias in the location of cell masks, cell positions were horizontally and vertically offset by 50% of the original dimensions and the analysis was repeated. The direction of microtubule growth was normalised for each embryo by dividing the events in each direction by the total number of events. The normalised directional ratio compares the upward versus downward EB1-GFP growth relatively to the dorsoventral axis of embryos and cellular elongation. *Statistical analysis*: Statistical analysis was performed in GraphPad Prism 8.0 (https://www.graphpad.com/scientific-software/prism/). The t-test (with Welch’s correction for data not normally distributed) or one-way ANOVA was used to compare two or more groups, respectively. The Smirnov-Kolmogorov test was used to compare differences in distributions of Arf1-GFP relative distances. We used a minimum sample size of 6 embryos per group. The specific details of each analysis are outlined in each figure legend. All graphs were made in either GraphPad Prism or R (ggplot package).

## Appendix E Monte Carlo parameter exploration

The full stochastic simulation of the motors on microtubules require fourteen parameters: the number of MTs on each side of the cell, the cell’s length, the parameters governing the MT dynamics, and the parameters governing the motor dynamics. In order to investigate the qualitative differences between the motor families’ absorption times without ignoring the uncertainty in all the parameter values, we perform a Monte Carlo simulation, which empirically estimates the first four moments of the kinesin and dynein arrival time distributions for randomly-sampled MT and motor parameter values. However, to focus on the walk-off effect, we chose to assign motors the same statistical characteristics (e.g. probability of picking up the back head and stepping forward, etc.), except for the preferred direction of travel.

At each iteration of the MC algorithm, the two absorption time distributions are estimated using 10^4^ realizations, with the discrete MT parameters uniformly sampled from the following ranges: cell size: *N*; ∈ {100, …, 1000}; and number of left- and right-directed MTs: *N*_*L*_, *N*_*R*_ ∈ {1, …, 20}. Under the assumption that the biologically-relevant MT parameter ranges, rescue *α*′ and catastrophe *β*′ rates are much smaller than the polymerization *α* and depolymerization *β* rates [58, 85], we fix the time unit so that the MT growth rate is *α* = 10^3^, and uniformly sample the other parameters from the ranges: *β* ∈ (2 · 10^3^, 6 · 10^3^), *α*′ ∈ (2, 10), *β*′ ∈ (*α*′*/*10, *α*′*/*2).

Generalizing [59] to the case of a system of finite antiparallel MTs, we introduce a hinged walker with three possible states: the fully attached state, in which both heads of the hinge are attached to adjacent MT dimers; the semi-attached state, in which one head is attached to a dimer, and the second head is free; and the diffusing state, in which both heads are detached. In the fully attached state, the back head detaches at a rate *β*_*b*_, and the front head detaches at a rate *β*_*f*_. From this semiat<u>ta</u>ched state, the walker either fully detaches at a rate 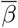, or the free head reattaches at the rate *ω*, with probability *p* of attaching in front of the attached head. In the detached state, the motor performs a random walk at a rate *η*, and reattaches (into the semiattached state) with uniform probability to any of the *N* available MTs at the rate *γN*, where *N* is the number of MTs reaching the motor’s position from either side of the bundle. This is motivated by the fact that reattachment is a chemical process. In the case a motor is attached to a depolymerizing MT, the motor switches to the detached state as soon as the dimer it is attached to disappears. Finally, once a motor arrives at either end of the MT, it is absorbed on the boundary, and no longer undergoes any dynamics.

Each pair of kinesin and dynein motors is initialized at the center of the cell. The kinesin stepping parameters are sampled uniformly from *p* ∈ (0, 1), *β*_*b*_, *β*_*f*_, *ω* ∈ (0, 2 ·10^4^), and are accepted if they satisfy

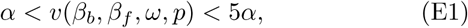

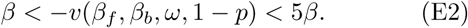

Inequality (E1) and (E2) enforce the conditions that kinesin outrun MT polymerization, and dynein depolymerization. Upon falloff, the motors diffuse in a lattice next to the MT bundle, until they re-attach to a MT. Since diffusion is known to be slower than active transport, we sample the diffusion rate from *η* ∈ (50, 500); and since motors have been observed to diffuse for some time before reattaching to a MT, we sample the reattachment rate from *γ* ∈ (*η/*10, *η/*2). Finally, because motors are likelier to step forwa<u>r</u>d than to fall of a MT, we sample the falloff rate from 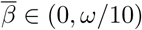.

The code for the stochastic simulations is available from the authors upon request.

## Appendix F MT length equilibrium distribution

Here we show that that the MT length equilibrium distribution is geometric. To find it a finite-length MT model in Fig.1a we modify the derivation in [86]. Consider the probabilities *p*_*n*_(*t*) and *q*_*n*_(*t*) that a MT of length *n* dimers is in a polymerizing or depolymerizing states, *P*_*n*_ or *D*_*n*_, respectively, and the maximum length of the MT is *N* + 1 dimers. Then *p*_*n*_(*t*) and *q*_*n*_(*t*) satisfy the following ODEs for the boundary dimers

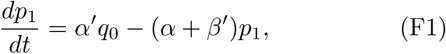

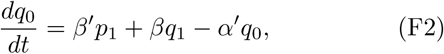

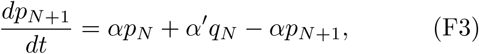

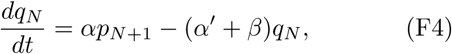

and for the internal dimers (1 ≤ *n* ≤ *N* − 1),

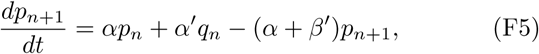

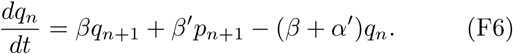

subject to the constraint

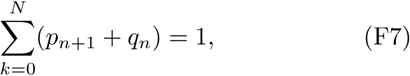

For 2 ≤ *n* ≤ *N*

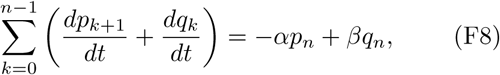

which in steady state reduces to

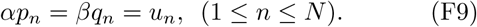

In view of (F5),(F4) and (F1), it leads to

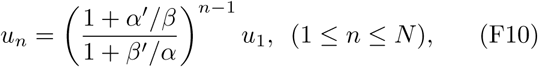

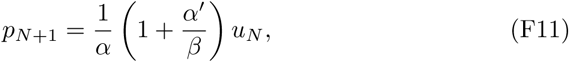

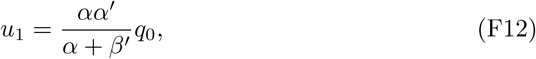

where *q*_0_ is chosen so that (F7) is satisfied.

## ACKNOWLEDGMENTS

This research was supported by the Leverhulme trust RPG-2017-249 (N.A.B, and L.C.); BBSRC BB/P007503/1 (N.A.B.); and the Royal Society of Edinburgh and the Scottish Government Personal Research Fellowship (L.C.).

## Author contributions

N.A.B. and L.C. designed the research and experiments; G.Z. performed stochastic simulations; G.Z. and L.C. developed the analytical model; V.A. and N.A.B. did biological experiments; G.Z., V.A, N.A.B, and L.C. wrote the manuscript. The authors declare no conflict of interest. N.A.B and L.C. contributed equally to this work.

